# Computational modeling of optogenetic control in human atrial tissue: The role of fibrosis type and illumination strategy

**DOI:** 10.1101/2025.09.26.678713

**Authors:** Heqing Zhan, Kaiqi Liu, Chuan’an Wei, Dongdong Deng, Yuanbo Yu

## Abstract

Atrial fibrosis promotes atrial fibrillation by stabilizing spiral waves, with focal fibrosis causing strong anchoring that resists termination. Optogenetics offers a low-energy alternative for rhythm control, but its efficacy in human atrial tissue remains under investigation. Here, we developed a two-dimensional computational model of human atrial tissue expressing the light-sensitive protein GtACR1, coupled with fibroblasts, to simulate spiral wave dynamics under varying degrees (0%–100%) and types (focal vs. diffuse) of fibrosis. We examined the effects of subthreshold illumination area (0%–100%) and spatial intensity profiles (uniform, parabolic, linear, exponential) on termination efficiency, and tested low-intensity periodic stimulation for wave drift induction. Results show that the minimum light area required for termination increases with fibrosis degree. At 25%–50% fibrosis, focal substrates require larger illumination areas than diffuse ones, confirming stronger anchoring. Crucially, the relationship between illumination area and termination time is non-monotonic. An “optimal light range” exists: smaller areas fail to eliminate the wave, while excessively large areas suppress wavefront collisions, prolonging termination. In diffuse fibrosis, low-intensity periodic illumination successfully induces drift and boundary collision, enabling self-termination. In contrast, in focal fibrosis, the spiral wave core remains anchored, and drift cannot be initiated by modulating stimulation frequency or intensity alone. Our findings demonstrate that fibrosis type and extent critically influence optogenetic control efficacy. The existence of an optimal illumination strategy highlights the need for spatially tailored interventions. These results provide a mechanistic basis for developing individualized, low-energy optogenetic defibrillation protocols.

**Author summary:** Atrial fibrillation, a common heart rhythm disorder, is often sustained by rotating electrical waves in the heart muscle. Scarring (fibrosis) in the atria can anchor these waves, making them harder to eliminate. Optogenetics — a technique that uses light to control genetically modified heart cells—offers a promising low-energy approach to stop these dangerous rhythms. However, how different patterns of scarring affect the success of optical interventions remains unclear.

In this study, we developed a computational model of human atrial tissue expressing a light-sensitive protein (GtACR1) to investigate how two major types of fibrosis—localized “focal” scars versus widespread “diffuse” scarring — influence the effectiveness of light-based rhythm control. We found that focal fibrosis strongly anchors spiral waves, requiring larger illuminated areas for termination, while diffuse fibrosis allows more efficient disruption with smaller light zones. Crucially, we discovered an “optimal light range”: too little light fails to stop the wave, but too much can paradoxically prolong termination by suppressing wavefront collisions.

Furthermore, we show that low-intensity periodic light can induce drift and self-termination in diffuse fibrotic tissue, but fails in focal substrates due to structural anchoring. Our findings highlight that one-size-fits-all optical strategies are suboptimal, and that personalized illumination protocols—tailored to individual fibrosis patterns—could enable safer, lower-energy defibrillation in the future.

## Introduction

Cardiovascular disease remains the leading cause of death worldwide, with cardiac arrhythmias playing a pivotal role in its morbidity and mortality[1-3]. Atrial fibrillation (AF), one of the most prevalent and debilitating persistent arrhythmias in clinical practice, is primarily sustained by self-perpetuating spiral waves in the atrial tissue[4, 5]. Atrial fibrosis, a key pathological substrate in the initiation and maintenance of AF, alters myocardial conduction properties and promotes the formation, stabilization, and anchoring of spiral waves, making it a critical factor in understanding the underlying mechanisms of AF[6, 7].

Clinically, electrical defibrillation is a highly effective method for terminating AF by delivering a high-energy shock to depolarize the entire heart. However, despite its high success rate, this approach is associated with significant drawbacks[8], including severe pain, myocardial trauma, and tissue damage, which limit its long-term applicability[9, 10]. Even low-energy implantable devices, though promising, have faced poor patient tolerance due to discomfort during cardioversion[11]. These limitations have driven the search for safer, low-energy, and more precise alternatives for rhythm control[12, 13].

In recent years, optogenetics has emerged as a transformative technology for cardiac rhythm modulation[14, 15]. By genetically expressing light-sensitive ion channels—such as the anion channelrhodopsin GtACR1—in cardiomyocytes, specific wavelengths of light can be used to precisely control membrane potential, inducing hyperpolarization or depolarization[16, 17]. Preclinical studies have demonstrated that subthreshold optical stimulation can effectively terminate life-threatening arrhythmias in small animal hearts, offering a promising pathway toward “optical defibrillation” with minimal energy consumption[18, 19]. Nevertheless, clinical translation in humans remains hindered by concerns regarding the safety of viral gene delivery, potential immunogenicity of exogenous opsins, and limitations in light delivery technology.

Compared to experimental studies, computational modeling offers distinct advantages. It enables precise, independent control over critical parameters such as fibrosis distribution, illumination intensity, spatial coverage, and temporal patterns, allowing for fine-grained investigation of spiral wave dynamics under highly reproducible and controllable conditions[20, 21]. Simulations eliminate the ethical constraints, high costs, and inter-subject variability inherent in animal experiments. Moreover, they permit virtual ablation, reverse engineering, and real-time visualization of transmembrane potentials and wavefront interactions—capabilities that facilitate mechanistic insights and guide the design of targeted interventions[22]. Thus, computational simulation serves not only as a vital surrogate in the current phase of optogenetic cardiac research but also as a powerful tool for advancing personalized and precision-based therapeutic strategies.

Despite these advances, existing optogenetic cardiac models often focus on homogeneous tissues or idealized obstacles, with insufficient attention to the clinically prevalent heterogeneity of atrial fibrosis—particularly the differential responses to optical control between focal and diffuse fibrotic patterns[23]. Focal fibrosis tends to create structural anchors that stabilize spiral waves through anchoring, whereas diffuse fibrosis increases conduction heterogeneity, promoting wave fragmentation and dynamic instability[24]. These distinct substrates may profoundly influence the efficacy of optogenetic interventions[25, 26]. Furthermore, prior studies have typically employed fixed illumination intensities or regular stimulation geometries, overlooking the nonlinear dynamical effects that may arise from spatial illumination patterns (e.g., area, shape) and low-intensity periodic stimulation, such as wave drift, boundary collision, or the existence of an “optimal light range[27].”

To address these gaps, we developed a two-dimensional (2D) computational model of human atrial tissue expressing GtACR1, systematically comparing the optogenetic control of spiral waves under varying degrees (0%, 25%, 50%, 75%, 100%) and types (focal vs. diffuse) of fibrosis. We specifically investigated the effects of illumination area (0%–100% of tissue) and intensity distribution patterns (uniform, parabolic, linearly increasing, exponentially increasing) on termination efficiency, and further analyzed the role of low-intensity periodic illumination in inducing spiral wave drift. This study aims to test the following hypotheses: (1) the degree and type of fibrosis jointly determine the minimum light area and time required for successful termination; (2) focal fibrosis, due to its strong anchoring effect, exhibits lower termination efficiency than diffuse fibrosis; and (3) an “optimal light range” exists—too small an area fails to extinguish the spiral wave, while too large an area may suppress wavefront collisions and prolong termination time. If validated, these findings will provide a mechanistic foundation for patient-specific optogenetic defibrillation strategies, paving the way for low-energy, high-precision interventions in the treatment of atrial fibrillation.

## Methods

A 2D computational model of human atrial tissue expressing the light-sensitive protein GtACR1 was developed to investigate spiral wave dynamics and their optogenetic control under different fibrosis patterns. The simulation program was implemented in Fortran. At the cellular level, all state variables were updated using the forward Euler method with a temporal resolution of 10 μ s to ensure numerical stability. At the tissue level, the reaction-diffusion equation was solved using the finite difference method with a five-point central difference scheme for spatial discretization of the Laplacian operator.

### Cellular-level model

The cellular model integrates three key components: (1) a human atrial myocyte model, (2) a fibroblast model, and (3) a dynamic model of the opsin GtACR1[28-30]. The time evolution of the transmembrane potential *V*_m_ of a single atrial myocyte is governed by:

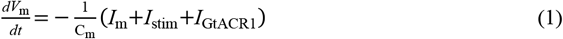

where *V*_m_ represents the atrial muscle cell membrane potential, C_m_ is the atrial muscle cell membrane capacitance, *I*_m_ is the sum of all transmembrane case currents, *I*_stim_ is the electrical stimulation current, and *I*_GtACR1_ is the GtACR1 current.

When an atrial myocyte is coupled to fibroblasts, the system of equations becomes:

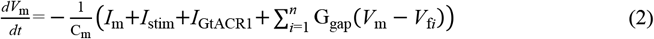

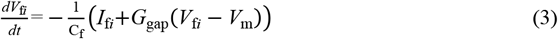

Where *V*_fi_ is the transmembrane potential of the *i*th coupled fibroblast, *I*_fi_ is the output current of the fibroblast, C_f_ is the fibroblast membrane capacitance, G_gap_ is the intercellular conductance, and *n* is the number of fibroblasts coupled to each myocyte. In this study, G_gap_ = 2.0 nS and *n* = 4 were fixed across all simulations to maintain model consistency, with the focus placed on the effects of illumination strategies.

### Tissue-level model and fibrosis representation

At the tissue level, electrical propagation is described by the monodomain reaction-diffusion equation[31]:

Non-fibrotic areas:

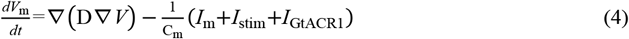

Fibrosis areas:

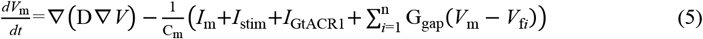

The simulation domain is a 101×101 square grid with a spatial resolution of 0.025 cm, corresponding to a physical size of approximately 2.525 cm × 2.525 cm. The isotropic diffusion coefficient is set to *D* = 0.00014 cm^2^/ms, yielding a baseline conduction velocity of approximately 43.9 cm/s.

Three distinct tissue configurations were considered in the simulations. The first configuration represented non-fibrotic tissue, in which all grid points were occupied by pure atrial myocytes. The second configuration modeled diffuse fibrosis, where fibroblast-coupled myocytes were randomly and uniformly distributed across the entire domain at specified densities of 25%, 50%, 75%, and 100%. The third configuration represented focal fibrosis, in which fibroblast-coupled cells were spatially concentrated within a quasi-square region centered in the domain, with the local coupling density corresponding to the designated fibrosis degree (25%, 50%, 75%, or 100%). To account for impaired electrical conduction in fibrotic regions, intercellular conductance was reduced depending on cell type adjacency: a 10% reduction was applied at connections between a pure myocyte and a fibroblast-coupled myocyte, while a 30% reduction was imposed between two adjacent fibroblast-coupled myocytes[32].

### Simulation protocol and stimulation scheme

The simulation consists of two sequential phases. In Phase I, a stable spiral wave was induced using an S1-S2 cross-field stimulation protocol. In Phase II, optogenetic intervention was applied to evaluate the efficacy of spiral wave termination. Once a stable spiral wave was established, subthreshold optical stimulation was delivered with varying parameters. The light intensity ranged from 0.004 to 1.0 μW/mm^2^, and the spatial coverage of illumination was varied from 0% to 100% of the total tissue area. Different spatial intensity profiles were tested, including uniform, parabolic, linearly increasing, and exponentially increasing distributions, to assess the impact of illumination pattern on termination efficiency. All simulations are conducted under no-flux boundary conditions. Spiral wave termination is defined as the complete disappearance of electrical activity across the entire domain for a duration exceeding 300 ms.

## Results

### Action potential of human atrial myocytes and fibroblasts in the presence of GtACR1 regulation

Fig. 1 illustrates the cellular-level responses of GtACR1-transduced human atrial myocytes and fibroblasts to optical stimulation. Fig. 1a shows the simulated action potentials of an atrial myocyte (red trace) and a coupled fibroblast (green trace) under uniform illumination applied between 500 and 1500 ms (indicated by the light-blue bar). In the dark phase (before 500 ms), both cells maintain their respective resting potentials. Upon illumination, both undergo a rapid shift in membrane potential, transitioning to a sustained depolarized state centered around −40 mV. The fibroblast also exhibits a similar steady-state shift, reflecting direct modulation by GtACR1-mediated ion flux. Fig. 1b presents the time course of the GtACR1-mediated transmembrane current. At light onset (500 ms), a brief initial inward current (approximately −0.5 pA) is observed, followed by a transient outward current (∼+0.35 pA), and then a stable inward current that persists at approximately −0.3 pA for the duration of illumination. The presence of this light-evoked current correlates temporally with the altered membrane dynamics in both cell types, confirming functional expression of the opsin and its capacity to influence cellular electrophysiology.

**Fig. 1.**
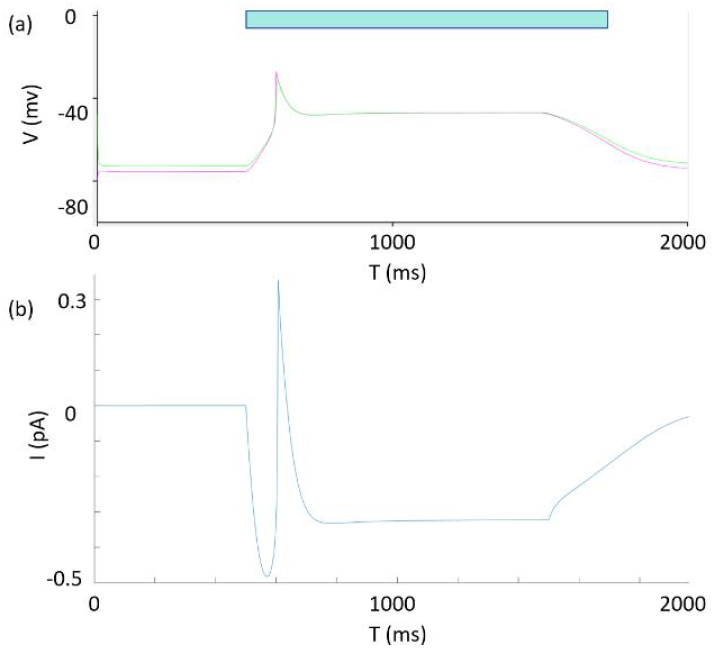
GtACR1-mediated electrophysiological responses in human atrial myocytes and fibroblasts. (a) Simulated action potentials of a GtACR1-expressing atrial myocyte (red) and a coupled fibroblast (green) during optical stimulation (light-blue bar: 500–1500 ms). (b) Time course of the GtACR1-mediated current, showing a triphasic response upon light activation.

### Effect of terminating spiral waves in different illumination ranges in different fibrosis conditions

Fig. 2 employed 2D computer simulations to investigate the termination efficacy of the light-sensitive protein GtACR1 on spiral waves in atrial myocytes with varying fibrosis degrees (0%, 25%, 50%, 75%, 100%) and distribution types (diffuse, focal) in different light ranges. At an illumination range of 35%, the results for pure atrial myocytes, 50% focal fibrosis (the fibrosis area is concentrated in the rectangular area at 50% center of the 2D plane), and 50% diffuse fibrosis are presented in Fig. 2.

**Fig. 2.**
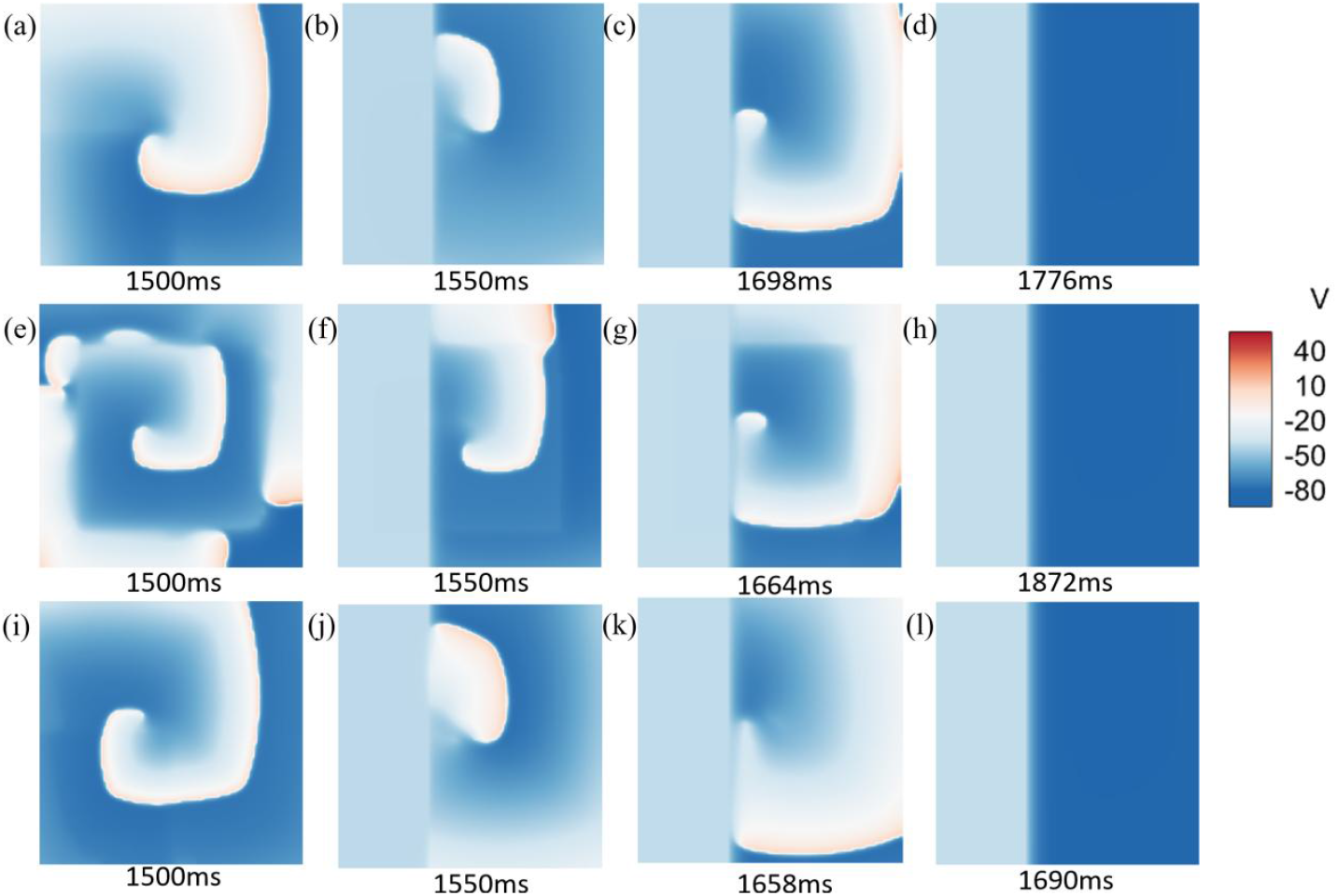
Spiral wave dissipation in atrial tissues under 35% light coverage (0.1 μW/mm^2^). (a-d) Spiral wave dissipation in pure atrial myocytes. (e-h) Spiral wave dissipation in focal atrial fibrosis (50% fibrosis). (i-l) Spiral wave dissipation in diffuse atrial fibrosis (50% fibrosis).

Fig. 2 illustrates the dissipation of spiral waves in a 2D plane under 35% light coverage across three conditions: pure atrial myocytes, focal atrial fibrosis (50% fibrosis), and diffuse atrial fibrosis (50% fibrosis). Specifically, in pure atrial myocytes, spiral waves completely dissipate in 276 ms (2a-2d); In focal atrial fibrosis (50% fibrosis), spiral waves take 372 ms to fully dissipate (e-h); In diffuse atrial fibrosis (50% fibrosis), spiral waves dissipate in 190 ms (i-l). This result contradicts the expectation that spiral waves in pure atrial myocytes would dissipate the fastest. To further investigate this discrepancy, additional simulations were conducted for pure atrial myocytes and the two fibrosis types (diffuse and focal) at varying degrees of fibrosis (0%, 25%, 50%, 75%, 100%) and different light coverage areas (0%–100% of the plane, increasing from left to right). The termination times of spiral waves under these conditions were calculated. The results were illustrated in Fig. 3.

**Fig. 3.**
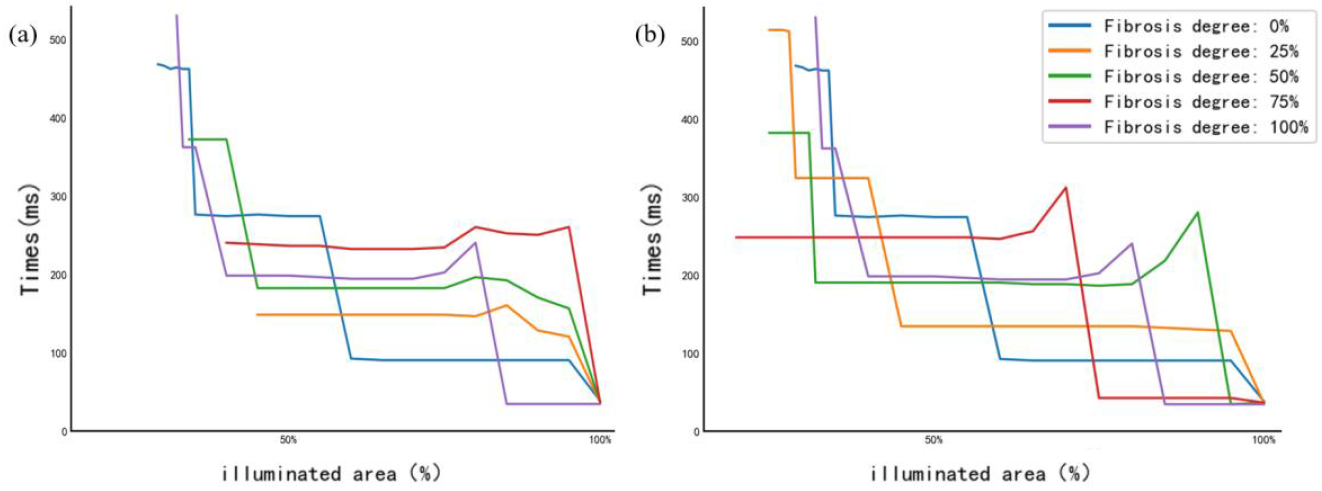
Relationship between spiral wave termination time and percentage of illuminated area in a 2D atrial tissue sheet under constant light intensity (0.1 μW/mm^2^). (a) focal fibrosis, (b) diffuse fibrosis. The x-axis represents the percentage of illuminated area, and the y-axis shows the time required for spiral wave termination. Different colored curves correspond to various degrees of fibrosis (0%, 25%, 50%, 75%, 100%).

In focal fibrosis conditions, the overall trend shows a decrease in spiral wave termination time as the illuminated area increases. For 25% fibrosis (orange line), measurable termination begins when illumination coverage exceeds 45%, with a termination time of 148 ms. Between 45% and 80% illumination, this time remains stable at approximately 145 ms before gradually decreasing further with increased illumination, ultimately reaching 36 ms under full coverage. In 75% fibrosis (red line), effective termination starts at 40% illumination (240 ms), maintaining around 240 ms within the 40%–75% range, then decreases to 36 ms at full illumination. For 100% fibrosis (purple line), there is an anomalous increase in termination time to 240 ms at 80% illumination, followed by a rapid decline to 34 ms at 85% illumination, stabilizing thereafter.

In diffuse fibrosis conditions, the overall trend also shows a reduction in termination time with increasing illuminated area. For 50% fibrosis (green line), termination time remains stable at approximately 188 ms across the 25%–80% illumination range; at 90% illumination, it anomalously rises to 280 ms before rapidly declining to stabilize at 36 ms under full coverage. For 75% fibrosis (red line), the termination time stays around 246 ms from 10% to 60% illumination, rising to 312 ms at 70% illumination before quickly dropping back to 36 ms at full illumination. In 100% fibrosis (purple line), the termination time ranges around 194 ms between 38% and 70% illumination, increasing to 240 ms at 80% illumination, then sharply decreasing to stabilize at 34 ms under full illumination.

Comparing the two types of fibrosis reveals that spiral waves in diffuse fibrotic substrates can be terminated with smaller illuminated areas compared to those in focal fibrosis. For example, the red curve (75% fibrosis) shows effective intervention starting at 10% illumination in the diffuse model, whereas in the focal model, effective termination requires at least 40% coverage.

What is more special is that although the overall relationship between the illumination area and the termination time was that the termination time decreases step by step as the illumination area increases, there were still some special cases where the termination time of the spiral wave increases with the illumination range. Here we take one example to explain it, as shown in Fig. 4.

**Fig. 4.**
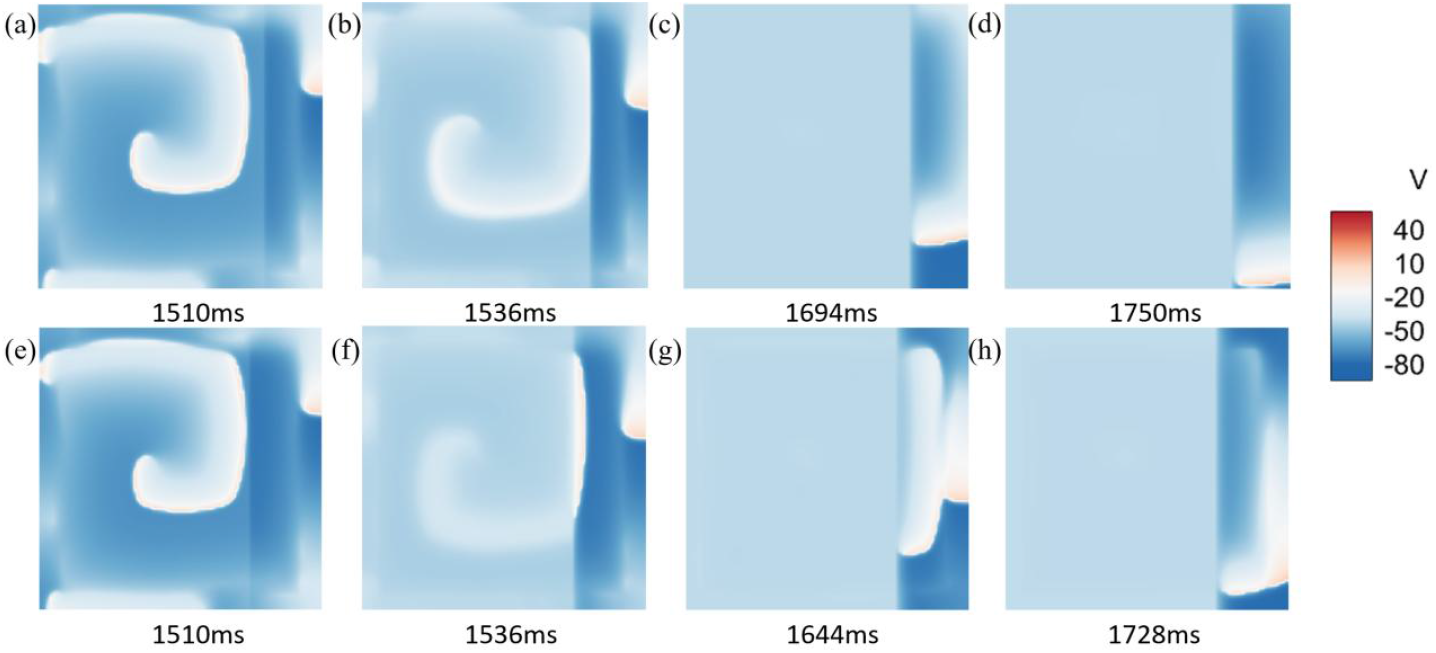
Collision-induced acceleration of spiral wave dissipation in focal fibrotic atrial tissue under suboptimal illumination (0.1 μW/mm^2^). (a-d) 80% illumination: complete suppression of the main spiral wave, leaving a slowly dissipating fragmented wave. (e-h) 75% illumination: residual wavefronts collide with fragmented waves, accelerating annihilation.

To investigate the phenomenon observed in Fig. 3—where spiral wave dissipation time paradoxically increases within a specific range of illumination coverage—we selected a representative case for simulation analysis. Fig. 4 presents dynamic snapshots of spiral wave evolution in a 75% fibrotic atrial tissue with focal fibrosis under 80% (a-d) and 75% (e-h) illumination coverage. It is observed that under 75% illumination, the spiral wave is not fully encompassed by the illuminated region; residual wavefronts persist outside the right boundary of the light-exposed area. These uncovered wavefronts continue to propagate and collide with fragmented wavelets remaining in the upper-right region of the tissue, subsequently accelerating propagation toward the lower-right corner, coalescing into a single wavefront, and ultimately dissipating. In contrast, under 80% illumination, the main body of the spiral wave is completely covered and rapidly suppressed, leaving only a small fragmented wave in the upper-right region to propagate into the non-illuminated zone before gradually dissipating. Compared to the 75% illumination condition, the 80% illumination scenario lacks wavefront fusion, and the dissipation time of the isolated fragmented wave is prolonged by approximately 26 ms.

### Influence of different light intensity distribution patterns on the termination effect of spiral waves

After investigating the effect of illumination coverage area on spiral wave termination, we further explored the influence of different light intensity distribution patterns on spiral wave dynamics in a 2D domain. Fig. 5 presents the evolution of spiral waves under four distinct light intensity profiles with full-field illumination coverage. The top row shows results for atrial tissue with diffuse fibrosis (a–d), and the bottom row for focal fibrosis (e–h), with both substrates having a fibrosis level of 50%. The four columns represent the following illumination patterns from left to right: (a, e) uniform illumination at 1 μW/mm^2^ across the entire domain; (b, f) parabolic illumination with peak intensity of 1 μW/mm^2^ at the center, tapering to 0 μW/mm^2^ at both lateral edges; (c, g) linearly increasing illumination from 0 μW/mm^2^ on the left edge to 1 μW/mm^2^ on the right edge; and (d, h) exponentially increasing illumination from 0 μW/mm^2^ on the left to 1 μW/mm^2^ on the right. Visually, no significant differences in spiral wave dynamics are observed across the four illumination patterns. The time required for spiral wave dissipation under each condition was recorded (see Table 1). The data show that, for both fibrotic substrates at the same fibrosis level, the termination times are nearly identical across all four illumination modes. This indicates that, under full-field illumination coverage, variations in the spatial distribution of light intensity do not significantly affect spiral wave termination.

**Table 1.** Termination duration of spiral waves under different illumination patterns, fibrosis levels, and tissue types (unit: ms)

| **Illumination Pattern** | Uniform illumination | Parabolic illumination | Linearly increasing illumination | Exponentially increasing illumination |
| --- | --- | --- | --- | --- |
| **Diffuse (0%)** | 28 | 28 | 28 | 28 |
| **Focal (0%)** | 28 | 28 | 28 | 28 |
| **Diffuse (25%)** | 26 | 26 | 26 | 26 |
| **Focal (25%)** | 28 | 28 | 28 | 28 |
| **Diffuse (50%)** | 28 | 28 | 28 | 28 |
| **Focal (50%)** | 28 | 30 | 28 | 28 |
| **Diffuse (75%)** | 28 | 28 | 28 | 28 |
| **Focal (75%)** | 28 | 28 | 30 | 28 |
| **Diffuse (100%)** | 26 | 26 | 26 | 26 |
| **Focal (100%)** | 26 | 26 | 26 | 26 |

**Fig. 5.**
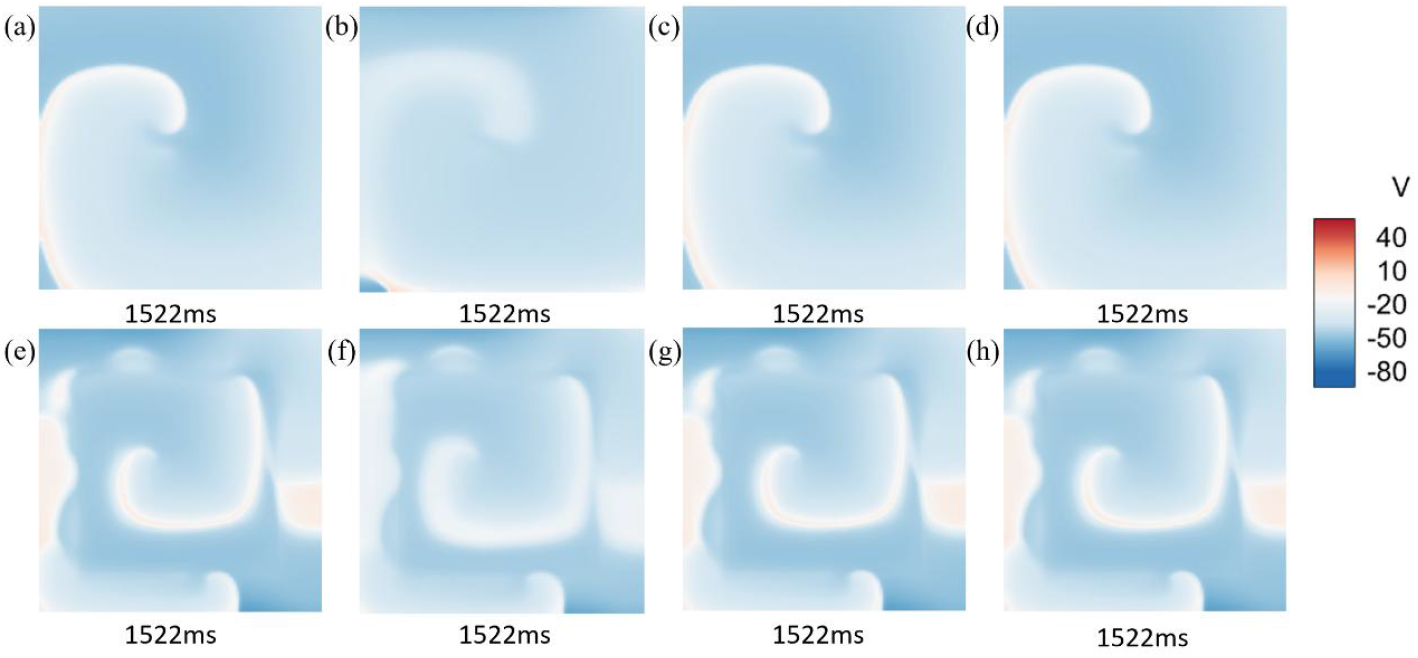
Effect of light intensity distribution on spiral wave dynamics in atrial tissue with diffuse and focal fibrosis under full-field illumination. (a, e) Uniform illumination (1 μW/mm^2^) on diffuse fibrosis (a) and focal fibrosis (e). (b, f) Parabolic illumination on diffuse fibrosis (b) and focal fibrosis (f). (c, g) Linearly increasing illumination from left to right on diffuse fibrosis (c) and focal fibrosis (g). (d, h) Exponentially increasing illumination from left to right on diffuse fibrosis (d) and focal fibrosis (h).

### Effect of low-intensity periodic light on spiral wave trajectory in fibrotic atrial tissue

Although we have previously established that a light intensity of 0.1 μW/mm^2^ is sufficient to terminate spiral waves in GtACR1-transduced atrial tissue with 50% fibrosis (Fig. 3), we further investigated the effects of periodic low-intensity global illumination to minimize potential photodamage. In this protocol, both the illumination duration and the dark interval were set to half the rotation period of the spiral wave (approximately 190 ms), resulting in repeated cycles of 95 ms of global uniform illumination followed by 95 ms of darkness. Fig. 6 illustrates the spiral wave dynamics under this illumination scheme in atrial tissue monolayers with diffuse fibrosis (a–d) and focal fibrosis (e–h), both at 50% fibrosis density. In diffuse fibrotic tissue, the spiral wave exhibits meandering motion: the wave tip initially drifts toward the upper-right corner, then shifts downward. In contrast, in focal fibrotic tissue, the spiral wave remains anchored near the center without significant drift. The wave tip trajectories under this periodic low-intensity illumination are shown in Fig. 7. In diffuse fibrosis (Fig. 7a), the wave tip moves from the center toward the upper-right region, then loops back to the left, approaches the center again, drifts upward once more, and finally propagates straight downward to the tissue boundary, where it annihilates. In focal fibrotic tissue (Fig. 7b), the wave tip remains confined to the central region with no notable displacement.

**Fig. 6.**
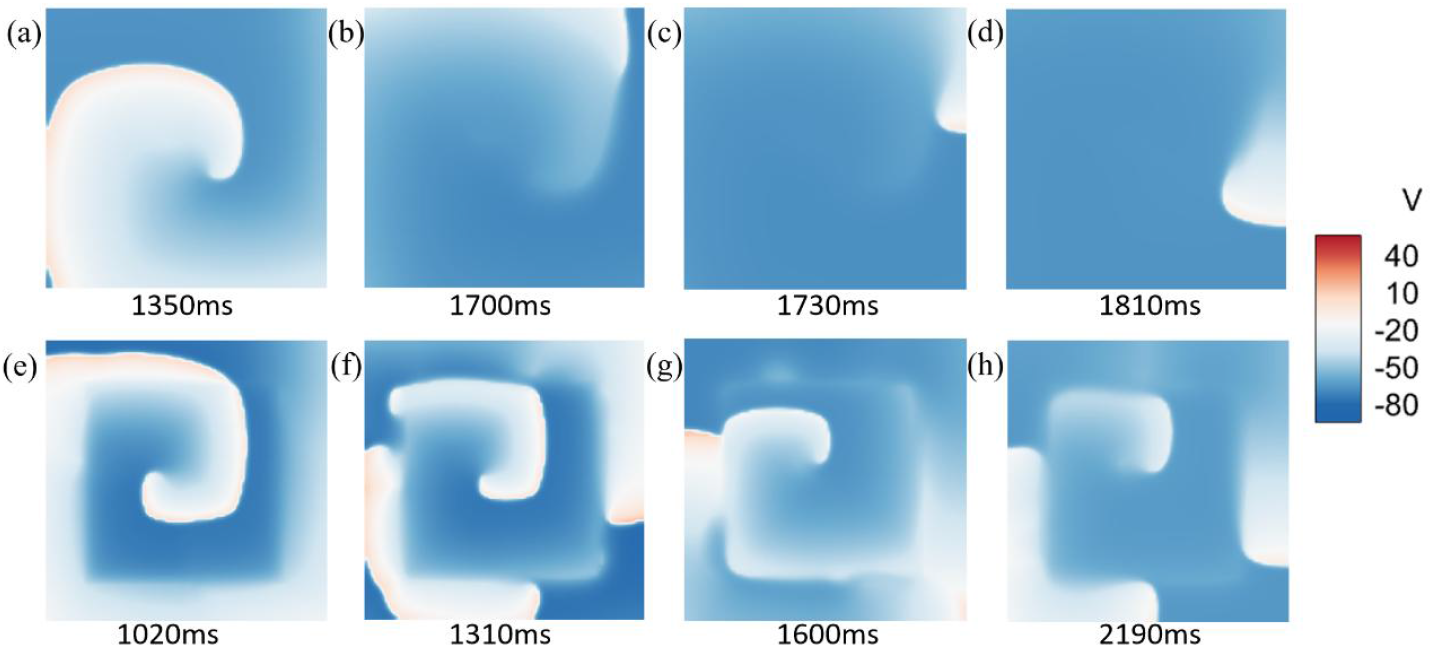
Spiral wave dynamics under periodic low-intensity illumination (0.004 μW/mm^2^) in atrial tissue with 50% diffuse and focal fibrosis. (a–d) Diffuse fibrosis: the spiral wave meanders, drifting first toward the upper-right and then downward. (e–h) Focal fibrosis: the spiral wave remains anchored near the center with no significant drift.

**Fig. 7.**
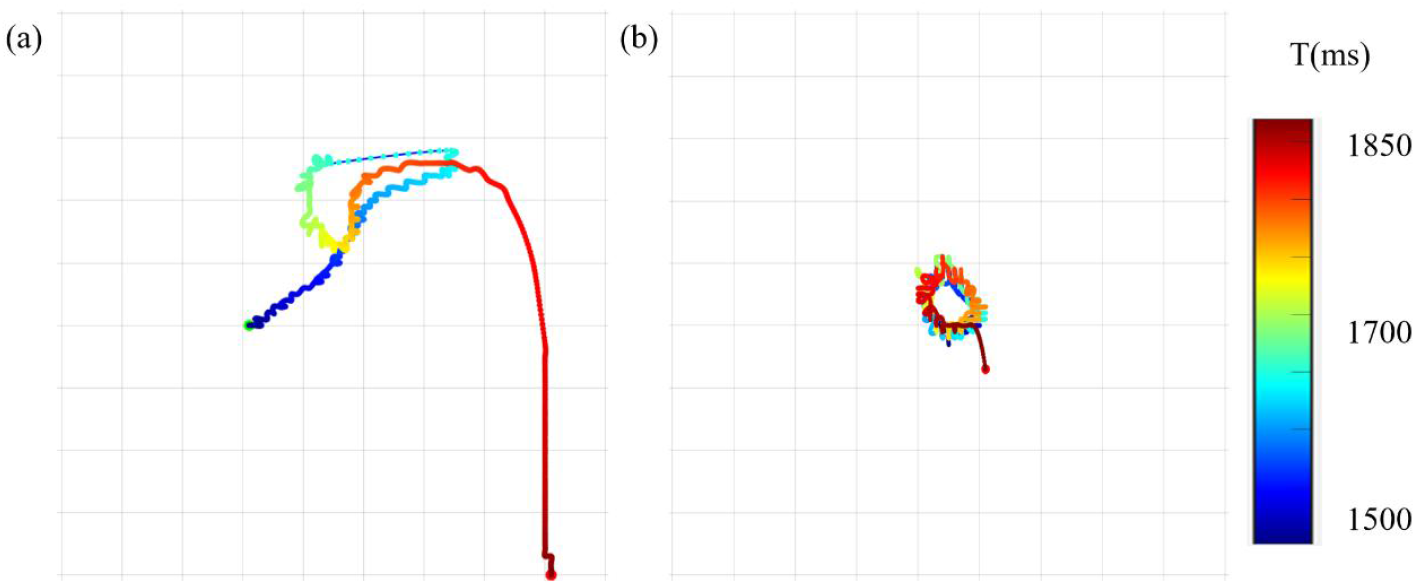
Trajectories of spiral wave tips under periodic low-intensity illumination (0.004 μW/mm^2^) in 50% fibrotic atrial tissue. (a) In diffuse fibrotic tissue, the wave tip follows a meandering path, eventually reaching the boundary and annihilating. (b) In focal fibrotic tissue, the wave tip remains confined to the central region without displacement.

## Discussion

This study aims to systematically investigate the mechanisms of optogenetic control of spiral wave dynamics in GtACR1-transduced atrial tissue, with a focus on how the extent and spatial distribution pattern (diffuse vs. focal) of atrial fibrosis influence the efficiency and pathways of optical termination. A 2D computational model based on human atrial myocyte electrophysiology was employed, incorporating coupling between atrial myocytes and fibroblasts and integrating the current dynamics of the light-sensitive protein GtACR1. By solving reaction-diffusion equations, the generation and evolution of spiral waves were simulated under varying degrees of fibrosis (0%, 25%, 50%, 75%, 100%) and distribution patterns (diffuse: randomly uniform; focal: centrally concentrated). On this basis, the effects of multiple illumination parameters on spiral wave termination were systematically evaluated, including illuminated area, light intensity (0.1 μW/mm^2^ for termination, 0.004 μW/mm^2^ for inducing drift), spatial intensity profiles (uniform, parabolic, linear/exponential gradients), and illumination timing (continuous vs. periodic pulsing). The results demonstrate that: (1) diffuse fibrosis is more sensitive to optical stimulation, enabling efficient spiral wave termination with smaller illuminated areas or low-intensity periodic illumination; (2) focal fibrosis, due to its strong structural anchoring effect, requires larger illumination coverage or high-intensity continuous intervention for effective termination; (3) there exists a non-monotonic relationship between illuminated area and termination time, with an “optimal range” beyond which excessive coverage may prolong termination by suppressing wave-wave interactions; (4) under full-field illumination, the spatial distribution of light intensity does not significantly affect termination efficiency; and (5) periodic low-intensity illumination can induce spiral wave drift and subsequent annihilation at tissue boundaries in diffuse fibrotic substrates, but fails to unanchor waves in focal fibrosis.

### The Critical Role of Fibrosis Spatial Pattern in Optogenetic Control Efficiency

In the field of optogenetic virtual heart modeling, various strategies have been developed to terminate spiral waves through light stimulation[33-35]. Studies have also demonstrated that in optogenetically modified cardiac tissue, regional illumination can induce directional linear drift of spiral waves[36, 37]. For instance, Hussaini et al. showed in a 2D model of adult mouse ventricles that localized illumination effectively controls the direction of spiral wave drift, which consistently moves toward the region of increasing light intensity[38]. However, these studies are predominantly based on healthy or homogeneous myocardial tissue models, with limited consideration of the structural heterogeneity associated with atrial fibrosis—a common clinical pathological substrate. Our study reveals the decisive influence of fibrosis spatial distribution (diffuse vs. focal) on the efficiency of optical control. Under 35% illumination coverage, spiral wave termination time is shortest in diffusely fibrotic tissue (190 ms), longest in focally fibrotic tissue (372 ms), and intermediate in non-fibrotic myocardium (276 ms). The underlying mechanism lies in the fact that diffuse fibrosis, characterized by randomly distributed fibroblasts, enhances global conduction heterogeneity, rendering the spiral wave electrophysiologically unstable and thus more susceptible to local optical perturbation. In contrast, focal fibrosis forms a high-density structural barrier in the central region, anchoring the spiral wave core and significantly enhancing its rotational stability, thereby increasing resistance to optogenetic intervention.

### Differential Thresholds for Effective Illumination in Diffuse and Focal Fibrotic Substrates

Although optogenetic induction of spiral wave drift has been demonstrated, systematic evaluation of the minimal effective illumination area across different fibrosis types remains lacking[38, 39]. In our simulations, spiral waves in diffusely fibrotic tissue exhibit higher sensitivity to low-coverage illumination, enabling effective termination with smaller illuminated areas, whereas focal fibrosis requires substantially larger coverage. For example, at 75% fibrosis, the diffuse model responds to intervention at just 10% illumination, while the focal model requires at least 40% coverage. This difference arises from distinct electrophysiological response mechanisms: in diffuse fibrosis, localized illumination generates steep repolarization gradients at the boundary between illuminated and non-illuminated regions, inducing conduction block or depolarization that disrupts the spiral wave structure. In focal fibrosis, however, the spiral wave core is surrounded by a dense cluster of fibrotic tissue, forming an electrophysiological “safe zone” that limits light penetration. Consequently, external illumination must be expanded to block the rotational pathway, necessitating broader spatial coverage for effective termination.

### Non-monotonic Dependence of Termination Time on Illuminated Area

Regarding control mechanisms, Ochs et al. demonstrated that the GtACR1 protein can terminate spiral waves under sub-threshold illumination, yet they did not investigate the complex relationship between illumination area and termination efficiency[28]. Our study reveals that while termination time generally decreases in a “stepwise” manner with increasing illuminated area, an anomalous non-monotonic effect occurs in specific regimes—termination time actually increases with larger illumination. For instance, in 75% diffuse fibrosis, termination time under 70% illumination exceeds that under 60% coverage. The fundamental mechanism lies in the competition between “suppression” and “interaction” in wave dynamics: moderate non-illuminated regions preserve residual wavefronts that can collide and merge with fragmented wavelets, accelerating annihilation. However, when illumination is excessive but not fully covering, the main spiral wave is strongly suppressed, leaving isolated wave fragments to decay slowly in non-illuminated zones, lacking effective wave-wave interactions and thus prolonging termination. This indicates the existence of an “optimal illumination range,” beyond which expanding coverage may counterproductively delay termination.

### Uniform Illumination Suffices for Global Termination

Furthermore, studies suggest that global high-intensity illumination may be required for optogenetic pacing to fill excitable gaps, highlighting an essential distinction in illumination requirements between different control objectives (e.g., pacing vs. defibrillation)[40]. In this study, focusing on defibrillation, we find that when full-field coverage is achieved, different spatial intensity profiles (e.g., uniform, parabolic, linear, or exponential gradients) have no significant impact on spiral wave termination efficiency. As long as the light intensity reaches the effective inhibition threshold (0.1 μW/mm^2^), the entire myocardial tissue synchronously enters a hyperpolarized state, blocking all excitable pathways. This indicates that termination efficiency primarily depends on whether the minimum effective intensity covers the entire domain, rather than on local intensity gradients. Therefore, under sub-threshold conditions, spatial variations in light intensity contribute negligibly to overall suppression, supporting the use of simple, uniform illumination in clinical applications.

### Structural Constraints on Spiral Wave Drift in Anchored Rotors

Hussaini et al. observed that localized illumination can guide spiral wave drift, but their model did not account for structural anchoring effects induced by fibrosis[38]. Our simulation results reveal distinct responses to periodic low-intensity illumination (0.004 μW/mm^2^, 95 ms on/95 ms off) in different fibrotic substrates: in diffuse fibrosis, the spiral wave exhibits significant drift and eventually annihilates at tissue boundaries; in focal fibrosis, the wave remains anchored to the fibrotic cluster and shows no substantial displacement. The mechanism lies in the differential response of anchored rotors to resonance: low-intensity periodic illumination can resonate with the intrinsic rotation frequency of the spiral wave, gradually pushing it outward. In diffuse substrates, where the tissue matrix is relatively uniform, drift resistance is low, allowing the wave core to migrate toward the boundary. In focal substrates, however, structural anchoring creates a strong “tethering effect” that firmly fixes the wave core, preventing effective drift. This demonstrates that structural anchoring poses a major barrier to low-intensity, non-invasive interventions. For such patients, sustained high-intensity illumination is required to directly eliminate the anchor. Our findings emphasize that clinical translation must adopt a “tailored” approach based on patient-specific fibrosis patterns, rather than a one-size-fits-all strategy, providing critical guidance for personalized optogenetic therapy.

### Limitations

This study advances the understanding of optogenetic control mechanisms of atrial spiral waves; however, several limitations warrant acknowledgment. First, simulations were conducted in (2D planar model, which does not incorporate the three-dimensional (3D) anatomical complexity of the real atria—such as the left atrial appendage, pulmonary vein ostia, and regional wall thickness variations—nor does it account for anisotropic conduction arising from myocardial fiber orientation. In 3D space, spiral waves may evolve into scroll waves, whose stability, drift dynamics, and response to optical stimulation may fundamentally differ from their 2D counterparts, thereby limiting the direct extrapolation of our findings to intact cardiac tissue. Second, fibrosis was modeled using idealized configurations, distinguishing only between diffuse (randomly and uniformly distributed) and focal (centrally concentrated) patterns. The model does not capture clinically observed features such as gradual fibrosis gradients, patchy distributions, or the structural organization of collagen networks, potentially underestimating the impact of complex microstructural heterogeneity on optogenetic control efficacy. Third, the model assumes uniform expression of the GtACR1 opsin across all cardiomyocytes and idealized, non-attenuating light propagation without scattering or absorption within the tissue. This neglects the optical attenuation inherent in biological tissues and potential heterogeneity in viral transduction efficiency, both of which may compromise spatial consistency in light responsiveness and affect the feasibility of low-intensity or localized intervention strategies. Finally, the illumination geometry was constrained to rectangular regions expanding from the left edge of the tissue, without systematically exploring more targeted illumination patterns—such as circular, annular, or dynamically tracked illumination centered on the spiral wave core—that could enable more efficient, energy-sparing ablation. While these simplifications facilitate mechanistic interpretation, future studies should aim to develop 3D multiscale models, integrate patient-specific fibrosis maps derived from clinical imaging, implement coupled opto-electrophysiological simulations incorporating light propagation physics, and explore adaptive illumination algorithms to enhance both physiological fidelity and translational potential.

## Conclusion

This computational study demonstrates that the spatial pattern of atrial fibrosis significantly influences optogenetic control of spiral waves. In GtACR1-transduced tissue, diffuse fibrosis is more sensitive to light intervention, allowing effective spiral wave termination with relatively small illuminated areas. In contrast, focal fibrosis anchors the spiral wave core, requiring larger illumination coverage for successful ablation. The relationship between illumination area and termination time is non-monotonic, with an “optimal range” beyond which larger areas may prolong termination—indicating that broader illumination is not always better. Global uniform illumination effectively terminates spiral waves, and the spatial profile of light intensity has minimal impact under full coverage. Low-intensity periodic illumination can induce drift and boundary collision in diffuse fibrosis, but fails in focal fibrosis due to structural anchoring. These findings suggest that optogenetic defibrillation strategies should be tailored to the specific fibrosis pattern. This work provides theoretical support for developing low-energy, precision-based approaches to arrhythmia control.

## Author contributions

**Conceptualization:** Heqing Zhan, Dongdong Deng

**Data curation:** Kaiqi Liu, Chuan’an Wei

**Formal analysis:** Heqing Zhan, Kaiqi Liu, Chuan’an Wei

**Funding acquisition:** Heqing Zhan, Dongdong Deng

**Methodology:** Heqing Zhan, Chuan’an Wei

**Project administration:** Yuanbo Yu

**Resources:** Heqing Zhan

**Software:** Kaiqi Liu

**Supervision:** Dongdong Deng, Yuanbo Yu

**Validation:** Heqing Zhan, Kaiqi Liu, Dongdong Deng, Yuanbo Yu

**Visualization:** Kaiqi Liu, Heqing Zhan

**Writing - original draft:** Heqing Zhan, Kaiqi Liu,

**Writing - review & editing:** Heqing Zhan, Dongdong Deng

## Data availability

Data will be made available on request.

## references

1. E.J. Benjamin, P. Muntner, A. Alonso, M.S. Bittencourt, C.W. Callaway, A.P. Carson, A.M. Chamberlain, A.R. Chang, S. Cheng, S.R. Das, et al. Heart Disease and Stroke Statistics-2019 Update: A Report From the American Heart Association. Circulation. 139 (10) (2019) e56–e528.

2. G.R. Mines. On dynamic equilibrium in the heart. J Physiol. 46 (4-5) (1913) 349–83.

3. D.J. Wilber, H. Garan, D. Finkelstein, E. Kelly, J. Newell, B. McGovern, and J.N. Ruskin. Out-of-hospital cardiac arrest. Use of electrophysiologic testing in the prediction of long-term outcome. N Engl J Med. 318 (1) (1988) 19–24.

4. V. Hakim and A. Karma. Theory of spiral wave dynamics in weakly excitable media: asymptotic reduction to a kinematic model and applications. Phys Rev E Stat Phys Plasmas Fluids Relat Interdiscip Topics. 60 (5 Pt A) (1999) 5073–105.

5. A.N. Zaikin and A.M. Zhabotinsky. Concentration wave propagation in two-dimensional liquid-phase self-oscillating system. Nature. 225 (5232) (1970) 535–7.

6. S. Nattel. Molecular and Cellular Mechanisms of Atrial Fibrosis in Atrial Fibrillation. JACC Clin Electrophysiol. 3 (5) (2017) 425–435.

7. C. Sohns and N.F. Marrouche. Atrial fibrillation and cardiac fibrosis. Eur Heart J. 41 (10) (2020) 1123–1131.

8. F. Godemann, C. Butter, F. Lampe, M. Linden, M. Schlegl, H.P. Schultheiss, and S. Behrens. Panic disorders and agoraphobia: side effects of treatment with an implantable cardioverter/defibrillator. Clin Cardiol. 27 (6) (2004) 321–6.

9. L.G. Tereshchenko, M.N. Faddis, B.J. Fetics, K.E. Zelik, I.R. Efimov, and R.D. Berger. Transient local injury current in right ventricular electrogram after implantable cardioverter-defibrillator shock predicts heart failure progression. J Am Coll Cardiol. 54 (9) (2009) 822–8.

10. G.M. Marcus, D.W. Chan, and R.F. Redberg. Recollection of pain due to inappropriate versus appropriate implantable cardioverter-defibrillator shocks. Pacing Clin Electrophysiol. 34 (3) (2011) 348–53.

11. J.C. Geller, S. Reek, C. Timmermans, T. Kayser, H.F. Tse, C. Wolpert, W. Jung, A.J. Camm, C.P. Lau, H.J. Wellens, et al. Treatment of atrial fibrillation with an implantable atrial defibrillator--long term results. Eur Heart J. 24 (23) (2003) 2083–9.

12. A.J. Moss, W.J. Hall, D.S. Cannom, J.P. Daubert, S.L. Higgins, H. Klein, J.H. Levine, S. Saksena, A.L. Waldo, D. Wilber, et al. Improved survival with an implanted defibrillator in patients with coronary disease at high risk for ventricular arrhythmia. Multicenter Automatic Defibrillator Implantation Trial Investigators. N Engl J Med. 335 (26) (1996) 1933–40.

13. S. Luther, F.H. Fenton, B.G. Kornreich, A. Squires, P. Bittihn, D. Hornung, M. Zabel, J. Flanders, A. Gladuli, L. Campoy, et al. Low-energy control of electrical turbulence in the heart. Nature. 475 (7355) (2011) 235–9.

14. A.B. Arrenberg, D.Y. Stainier, H. Baier, and J. Huisken. Optogenetic control of cardiac function. Science. 330 (6006) (2010) 971–4.

15. T. Bruegmann, D. Malan, M. Hesse, T. Beiert, C.J. Fuegemann, B.K. Fleischmann, and P. Sasse. Optogenetic control of heart muscle in vitro and in vivo. Nat Methods. 7 (11) (2010) 897–900.

16. R.A. Kopton, J.S. Baillie, S.A. Rafferty, R. Moss, C.M. Zgierski-Johnston, S.V. Prykhozhij, M.R. Stoyek, F.M. Smith, P. Kohl, T.A. Quinn, et al. Cardiac Electrophysiological Effects of Light-Activated Chloride Channels. Front Physiol. 9 (2018) 1806.

17. O.A. Sineshchekov, E.G. Govorunova, H. Li, and J.L. Spudich. Gating mechanisms of a natural anion channelrhodopsin. Proc Natl Acad Sci U S A. 112 (46) (2015) 14236–41.

18. W. Li, C.M. Ripplinger, Q. Lou, and I.R. Efimov. Multiple monophasic shocks improve electrotherapy of ventricular tachycardia in a rabbit model of chronic infarction. Heart Rhythm. 6 (7) (2009) 1020–7.

19. C.M. Ambrosi, C.M. Ripplinger, I.R. Efimov, and V.V. Fedorov. Termination of sustained atrial flutter and fibrillation using low-voltage multiple-shock therapy. Heart Rhythm. 8 (1) (2011) 101–8.

20. R. Majumder, I. Feola, A.S. Teplenin, A.A. de Vries, A.V. Panfilov, and D.A. Pijnappels. Optogenetics enables real-time spatiotemporal control over spiral wave dynamics in an excitable cardiac system. Elife. 7 (2018).

21. V. Biasci, L. Santini, G.A. Marchal, S. Hussaini, C. Ferrantini, R. Coppini, L.M. Loew, S. Luther, M. Campione, C. Poggesi, et al. Optogenetic manipulation of cardiac electrical dynamics using sub-threshold illumination: dissecting the role of cardiac alternans in terminating rapid rhythms. Basic Res Cardiol. 117 (1) (2022) 25.

22. T. Bruegmann, P.M. Boyle, C.C. Vogt, T.V. Karathanos, H.J. Arevalo, B.K. Fleischmann, N.A. Trayanova, and P. Sasse. Optogenetic defibrillation terminates ventricular arrhythmia in mouse hearts and human simulations. J Clin Invest. 126 (10) (2016) 3894–3904.

23. A. Defauw, P. Dawyndt, and A.V. Panfilov. Initiation and dynamics of a spiral wave around an ionic heterogeneity in a model for human cardiac tissue. Phys Rev E Stat Nonlin Soft Matter Phys. 88 (6) (2013) 062703.

24. A. Pumir, S. Sinha, S. Sridhar, M. Argentina, M. Hörning, S. Filippi, C. Cherubini, S. Luther, and V. Krinsky. Wave-train-induced termination of weakly anchored vortices in excitable media. Phys Rev E Stat Nonlin Soft Matter Phys. 81 (1 Pt 1) (2010) 010901.

25. G. Gottwald, A. Pumir, and V. Krinsky. Spiral wave drift induced by stimulating wave trains. Chaos. 11 (3) (2001) 487–494.

26. S.R. Kharche, I.V. Biktasheva, G. Seemann, H. Zhang, and V.N. Biktashev. A Computer Simulation Study of Anatomy Induced Drift of Spiral Waves in the Human Atrium. Biomed Res Int. 2015 (2015) 731386.

27. R.A. Quiñonez Uribe, S. Luther, L. Diaz-Maue, and C. Richter. Energy-Reduced Arrhythmia Termination Using Global Photostimulation in Optogenetic Murine Hearts. Front Physiol. 9 (2018) 1651.

28. A.R. Ochs, T.V. Karathanos, N.A. Trayanova, and P.M. Boyle. Optogenetic Stimulation Using Anion Channelrhodopsin (GtACR1) Facilitates Termination of Reentrant Arrhythmias With Low Light Energy Requirements: A Computational Study. Front Physiol. 12 (2021) 718622.

29. M. Courtemanche, R.J. Ramirez, and S. Nattel. Ionic mechanisms underlying human atrial action potential properties: insights from a mathematical model. Am J Physiol. 275 (1) (1998) H301–21.

30. D.E. Krummen, J.D. Bayer, J. Ho, G. Ho, M.R. Smetak, P. Clopton, N.A. Trayanova, and S.M. Narayan. Mechanisms of human atrial fibrillation initiation: clinical and computational studies of repolarization restitution and activation latency. Circ Arrhythm Electrophysiol. 5 (6) (2012) 1149–59.

31. M.M. Maleckar, J.L. Greenstein, W.R. Giles, and N.A. Trayanova. Electrotonic coupling between human atrial myocytes and fibroblasts alters myocyte excitability and repolarization. Biophys J. 97 (8) (2009) 2179–90.

32. K.A. MacCannell, H. Bazzazi, L. Chilton, Y. Shibukawa, R.B. Clark, and W.R. Giles. A mathematical model of electrotonic interactions between ventricular myocytes and fibroblasts. Biophys J. 92 (11) (2007) 4121–32.

33. P. Sasse, M. Funken, T. Beiert, and T. Bruegmann. Optogenetic Termination of Cardiac Arrhythmia: Mechanistic Enlightenment and Therapeutic Application? Front Physiol. 10 (2019) 675.

34. N. DeTal, A. Kaboudian, and F.H. Fenton. Terminating spiral waves with a single designed stimulus: Teleportation as the mechanism for defibrillation. Proc Natl Acad Sci U S A. 119 (24) (2022) e2117568119.

35. Q.H. Li, Y.X. Xia, S.X. Xu, Z. Song, J.T. Pan, A.V. Panfilov, and H. Zhang. Control of spiral waves in optogenetically modified cardiac tissue by periodic optical stimulation. Phys Rev E. 105 (4-1) (2022) 044210.

36. A.A. Nizamieva, I.Y. Kalita, M.M. Slotvitsky, A.K. Berezhnoy, N.S. Shubina, S.R. Frolova, V.A. Tsvelaya, and K.I. Agladze. Conduction of excitation waves and reentry drift on cardiac tissue with simulated photocontrol-varied excitability. Chaos. 33 (2) (2023) 023112.

37. Y.X. Xia, L.H. Xie, Y.J. He, J.T. Pan, A.V. Panfilov, and H. Zhang. Numerical study of the drift of scroll waves by optical feedback in cardiac tissue. Phys Rev E. 108 (6-1) (2023) 064406.

38. S. Hussaini, V. Venkatesan, V. Biasci, J.M. Romero Sepúlveda, R.A. Quiñonez Uribe, L. Sacconi, G. Bub, C. Richter, V. Krinski, U. Parlitz, et al. Drift and termination of spiral waves in optogenetically modified cardiac tissue at sub-threshold illumination. Elife. 10 (2021).

39. S. Hussaini, R. Majumder, V. Krinski, and S. Luther. In silico optical modulation of spiral wave trajectories in cardiac tissue. Pflugers Arch. 475 (12) (2023) 1453–1461.

40. T. Bruegmann, T. Beiert, C.C. Vogt, J.W. Schrickel, and P. Sasse. Optogenetic termination of atrial fibrillation in mice. Cardiovasc Res. 114 (5) (2018) 713–723.

